# Integration of spatial multiplexed protein imaging and transcriptomics in the human kidney tracks the regenerative potential timeline of proximal tubules

**DOI:** 10.1101/2024.11.26.625544

**Authors:** Mahla Asghari, Angela R. Sabo, Daria Barwinska, Ricardo Melo Ferreira, Michael Ferkowicz, William S. Bowen, Ying-Hua Cheng, Debora L. Gisch, Connor Gulbronson, Carrie L Phillips, Katherine J. Kelly, Timothy A. Sutton, James C Williams, Miguel Vazquez, John O’Toole, Paul Palevsky, Sylvia E. Rosas, Sushrut S. Waikar, Krzysztof Kiryluk, Chirag Parikh, Jeff Hodgins, Pinaki Sarder, Ian H. De Boer, Jonathan Himmelfarb, Matthias Kretzler, Kidney Precision Medicine Project, Sanjay Jain, Michael T. Eadon, Seth Winfree, Tarek M. El-Achkar, Pierre C Dagher

## Abstract

The organizational principles of nephronal segments are based on longstanding anatomical and physiological attributes that are closely linked to the homeostatic functions of the kidney. Novel molecular approaches have recently uncovered layers of deeper signatures and states in tubular cells that arise at various timepoints on the spectrum between health and disease. For example, a dedifferentiated state of proximal tubular cells with mesenchymal stemness markers is frequently seen after injury. The persistence of such a state is associated with failed repair. Here, we introduce a novel analytical pipeline applied to highly multiplexed spatial protein imaging to characterize proximal tubular subpopulations and neighborhoods in reference and disease human kidney tissue. The results were validated and extended through integration with spatial and single cell transcriptomics. We demonstrate that, in reference tissue, a large proportion of S1 and S2 proximal tubular epithelial cells express THY1, a mesenchymal stromal and stem cell marker that regulates differentiation. Kidney disease is associated with loss of THY1 and transition towards expression of PROM1, another stem cell marker shown recently to be linked to failed repair. We demonstrate that the trajectory of proximal tubular cells to THY1 expression is clearly distinct from that of PROM1, and that a state with PROM1 expression is associated with niches of inflammation. Our data support a model in which the interplay between THY1 and PROM1 expression in proximal tubules associates with their regenerative potential and marks the timeline of disease progression.

## Introduction

The kidney maintains body homeostasis through the integrated functions of nephronal segments and numerous cell populations. High-resolution molecular interrogation techniques such as single cell RNA sequencing have underscored the complexity of the kidney, which is composed of at least 70 interacting cell types ^1-4^. Our understanding of these specialized cells has increased in granularity, since we now recognize the presence of subpopulations in what was thought to be uniform cell types. Furthermore, within a given population, recent depictions of various cell states that correlate with disease trajectory or outcomes usher new opportunities to uncover key pathways in the pathogenesis of kidney disease and precise biomarkers that clock the disease timeline^1,2^. In fact, some altered cells states (typically defined by the expression profile of specific stress or mesenchymal-associated genes or proteins) are associated with maladaptive or failed repair following injury and may predict the progression to chronic kidney disease^1^. Importantly, the power of single cell methodologies can be greatly expanded when analyzed in a “spatial” context. Indeed, the interaction of various cells in spatially defined microenvironments within the kidney likely governs many of these pathological processes^1,2,5-7^. Novel large scale multiplexed spatial protein imaging techniques such as CO-Detection by indEXing (CODEX) capture cell types and their distribution within the complex landscape of kidney tissue^7^. Applying such a spatially anchored analytical pipeline can then provide “single cell data” with spatial protein expression features. The power of such spatial protein-based analysis can be further augmented when integrated with spatial gene expression platforms such as spatial transcriptomics (ST), performed on a sequential section from the same specimen^4,8^. Such spatially anchored integrative analysis bridging protein to RNA at the single cell level will greatly improve our understanding of the pathophysiology of kidney disease and the drivers of its progression.

As the most upstream tubular segment of the nephron, the proximal tubule is central to the functioning of the kidney and orchestrates tight communication with more distal segments^9,10^. It is also a frequent target of disease, and its pathology rapidly spreads to other cell populations. Many proximal tubular (PT) cell subpopulations have been described along the spectrum from health to disease^1,11-15^. While some are permanent components of the PT, others have a more fleeting existence, appearing or vanishing at specific points in the timeline of disease. Of particular interest are subtypes of PTs that undergo maladaptive repair, ushering the progressing towards fibrosis^11-13^. These cells are defined by the persistent expression of several injury markers such as *HAVCR1* (KIM1) or *VCAM1*^13,14,16^. These PT subtypes have been well characterized in both experimental models of kidney injury and in human kidney tissue from patients with kidney disease^1,11-13^. In addition, genes that typically mark stem cells and are involved in repair, such as *PROM1* (CD133) and *SOX 9*, are upregulated post injury^17-19^. However, persistent activation of these genes is frequently associated with maladaptive repair and progressive fibrosis^1,18,20^.

Thymus antigen 1 (THY1, also known as CD90) is a small glycoprotein that was originally described as a T cell marker in the thymus, but later found to be expressed in stem cells and a variety of neuronal, mesenchymal, stromal and endothelial cells^21^. THY1 interacts with various ligands to regulate signaling pathways related to cell growth and regeneration. For example, THY1 is thought to play a key regulatory role in the differentiation of mesenchymal stromal cells^21,22^. A recent report by Wu and colleagues showed that THY1 is expressed in human proximal tubules^23^. The function of THY1 in the kidney is unclear and its expression in various PT subpopulations has not been investigated. Because of its association with stem cells and regeneration, there has been a growing interest in its potential role in kidney development and kidney repair after injury^21,23^. THY1 also exists in a soluble form, and an inverse correlation between soluble THY1 levels and kidney function has been described^21,23^. Interestingly, despite reports suggesting THY1 expression in mouse kidneys^21^, previous data^24^ and various murine single cell expression databases^2,13,25^ have consistently demonstrated that THY1 is not expressed in mouse tubular cells. THY1 expression in the mouse kidneys is likely to originate from extra-tubular sources such as infiltrating immune cells. Whether THY1 expression in the kidney tubules is unique to humans remains unknown.

In this work, we aimed to leverage a CODEX analytical pipeline integrated with spatial and single cell transcriptomics to establish the presence and spatial location of distinct populations of THY1-positive and PROM1-positive PT cells. We also define the molecular trajectory and significance of these cells in human reference and disease kidney tissue. Our findings highlight the importance of PT cell states in understanding the timeline of kidney disease progression and suggest that THY1 loss and a shift towards high PROM1 states are associated with chronic kidney disease.

## Results

### Building a multiplexed imaging and analytical pipeline to decipher cell types in health and disease

To delineate the various cell types and cell states using spatial protein expression information, we first defined the identity of each cell in a set of healthy and diseased kidney tissue samples using a CODEX marker panel with structural, immune and injury markers (Supplemental Tables 1 & 2 and Figure S1). Summary demographics and diagnosis of reference and disease kidney tissue specimens are presented in Supplemental Table 1. The initial core maker panel applied to all specimens (23 markers) was subsequently extended to include more specialized markers (total 38) (see Supplemental Tables 1 & 2 for details by specimen). CODEX imaging was performed at large scale, scanning the entire tissue sections ^7^. Quality control steps, technical and biological reproducibility were described in the methods and shown in Supplemental Figure S2. Applying our imaging and analysis pipeline within a single analytical space using the core markers, we classified a total of 169,802 cells (Figure1a-1c). This approach was successful in identifying all major expected cell types in the kidney based on the protein expression profile including glomerular, proximal tubular, distal tubular, endothelial and immune cells. The cell classes were spatially mapped to the images to validate the cell labels (Figure 1d-1k).

**Figure 1:**
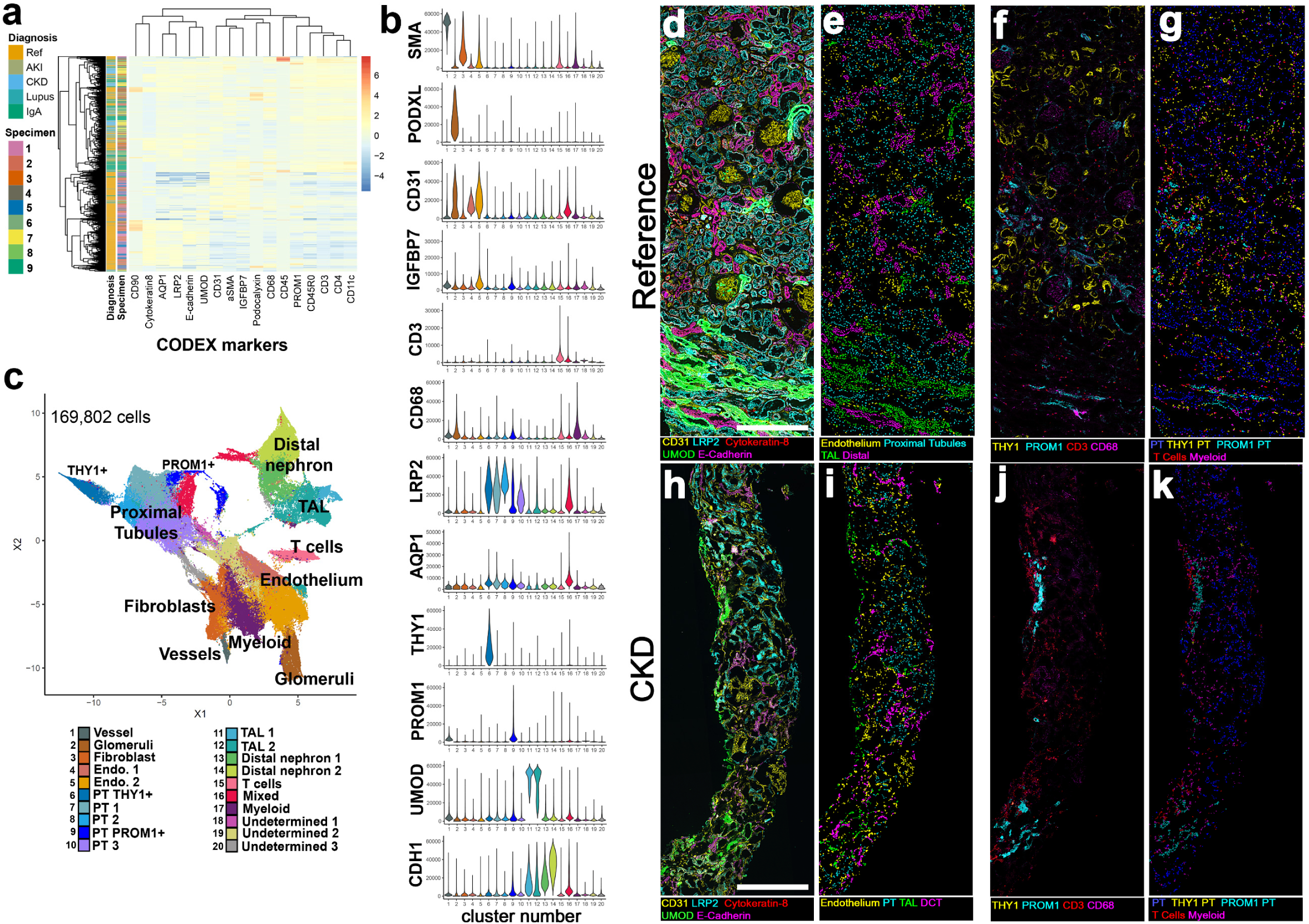
Cell classification based on CODEX imaging in reference and disease specimen. Reference and disease specimens were stained and imaged using CODEX. (a) Heatmap showing the z-scaled mean fluorescence intensity for CODEX markers for all the cells segmented from the images of reference and disease kidney tissue sections. The specimen identification number and diagnosis are indicated on the left. (b) Violin plots showing the mean fluorescence intensity distribution per cell of the markers indicated in each of the cell clusters. (c) UMAP plot of the Louvain clusters identified from the reference and disease specimens. Cell annotation was based on the marker expression profile (b), and from mapping back of the clusters to the images (d-k). Distal convoluted tubules, connecting segments and collecting ducts cells could not be individually resolved based on the markers used, and are labeled collectively as distal nephron. (d-g) Representative CODEX images of select markers in reference tissue (d & f) and the corresponding cell clusters mapped back as nuclear overlays (e & g). Scale bars = 500 μm. (h-k) Representative CODEX images of select markers in a kidney biopsy from CKD (h & j) and the corresponding cell clusters mapped back as nuclear overlays (i & k).

### Identifying PT cell sub-types and states using protein expression analysis with CODEX

To increase the resolution of PT cell subtypes, we re-clustered PT cells in a new analytical space (Figure 2). Clustering was predominantly based on the expression of LRP2, AQP1, and the presence or absence of CD68, THY1, PROM1 and IGFBP7(Figure 2a, b). We uncovered 2 PT subtypes that uniquely expressed THY1 or PROM1 (Figure 2a-d). As discussed, THY1 (also known as CD90) is a stem cell marker implicated in cell differentiation and regeneration^21^. PROM-1 is a marker of progenitor cells that is also expressed in cells undergoing adaptive or maladaptive repair^1,19^. The proximal tubular identity of these subtypes was confirmed morphologically by back-mapping the cell clusters onto the images (Figure 2 e-j). An extended CODEX panel applied to a disease tissue was also used to validate the profile of the PT subtypes, showing a protein expression signature consistent with injury (high KIM-1, p-cJUN and pMLKL) for PROM1-positive PTs as compared to THY1-positive or other PTs (Figure S3).

**Figure 2:**
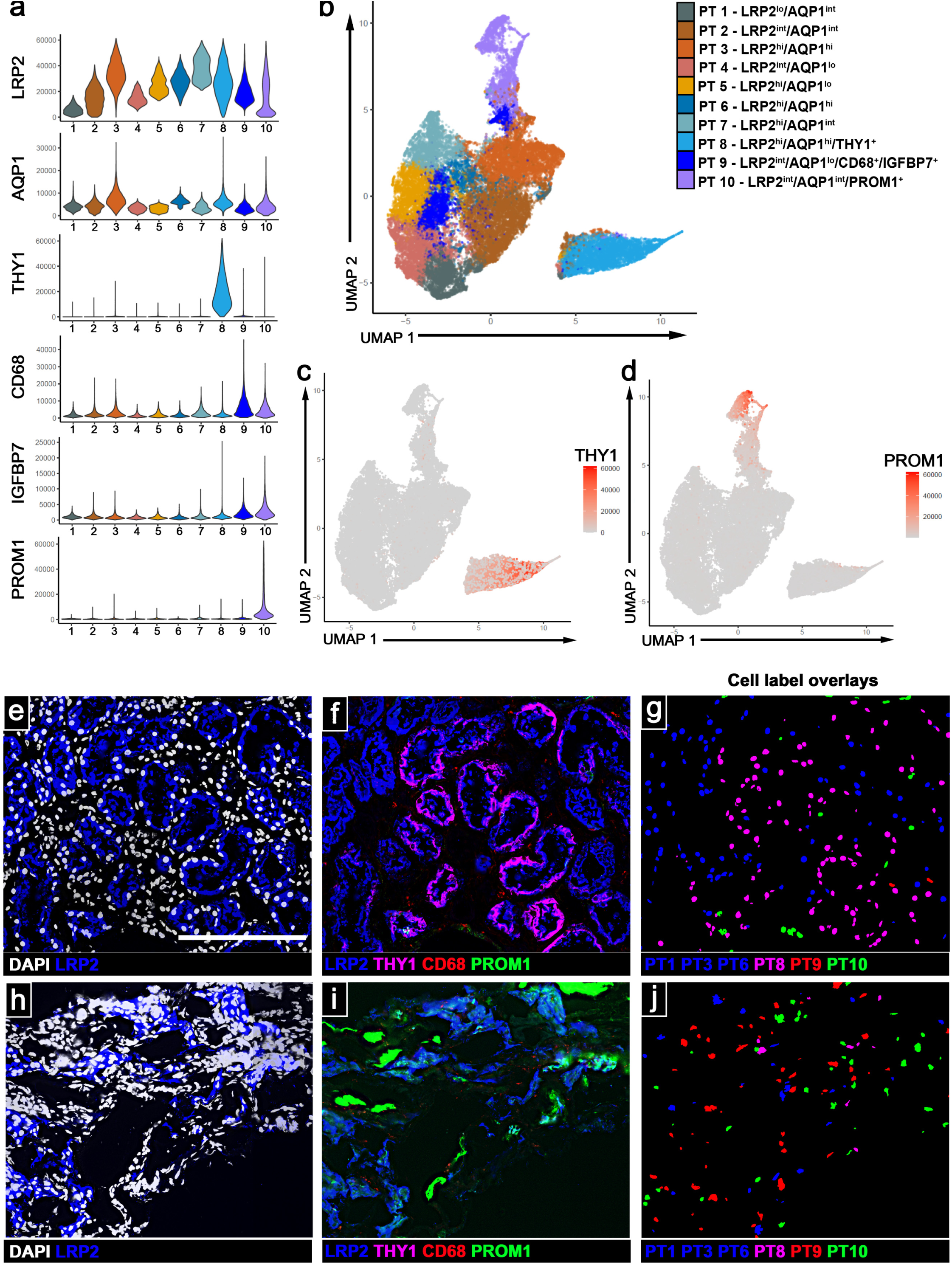
Sub-clustering of proximal tubular cells identifies PT groups expressing injury and regenerative state markers. Post CODEX and cell-type classification, proximal tubule cells were reclustered with a reduced set of molecular markers. (a) Violon plots showing the mean intensity distribution per cell of the markers relevant for each of the PT cell clusters. (b) UMAP plot visualizing the various PT clusters, highlighting the unique separation of the THY1-positive and PROM1-positive clusters. (c,d) Feature plots of THY1 and PROM1 expression. (e-g) representative images from reference showing staining of DAPI, LRP2, THY1, CD68, PROM1 (e,f) and mapping of the corresponding cell clusters as nuclear overlays (g). (h-j) representative images from disease tissue with similar staining (h,i) and mapping of the corresponding cell clusters (j). Scale bar = 200 μm

### Bridging THY1 expression from protein to RNA by integrating CODEX and spatial transcriptomics (ST)

PROM1 expression has been described in a subpopulation of PTs but the expression of THY1 in the human kidney and within PT cells is less well characterized. Here, we aimed to spatially define the relationship between THY1 protein and RNA expression. To determine the concordance of THY1 protein and mRNA expression, we used spatially co-registered sequential sections of human kidney cortex that underwent CODEX and ST (Figure 3a-c). ST spots overlaying THY1-positive PTs (defined as THY1 -positive, LRP2 positive by CODEX) were identified (Figure 3d-f) and found to have high *THY1* RNA by differential gene expression as compared to spots overlaying THY1*-*negative PT cells as defined by CODEX (Figure 3g). Additional genes were differentially upregulated in THY1-positive PT spots, including *MT1G, GATM, GPX3, APOE, PDZK1IP1, ALDOB* (Figure 3g). To validate and extend the findings at the single cell level, we used the single nuclear RNA sequencing (snRNA-seq) KPMP atlas to subset PTs (Figure 3h). We identified a PT subcluster (Cluster 5, arrow in Figure 3i) with a high-level of *THY1* expression, showing enrichment with similar genes to those uncovered by ST in CODEX-defined THY1-positive PTs (Figure 3g and 3i). This was also independently confirmed by comparing *THY1*-positive PTs gene expression to all other PTs in the snRNA-seq atlas (Figure 3j-3k). Pathway analysis with the DEGs in *THY1*-positive PTs revealed pathways related to cell proliferation and differentiation (Figure 3l), which are consistent with proposed functions of THY1 and its link to stemness^26^. We also confirmed distinct PT clusters with high *PROM1* expression (clusters 1 & 8 in Figure 3i), analogous to the findings with CODEX.

**Figure 3:**
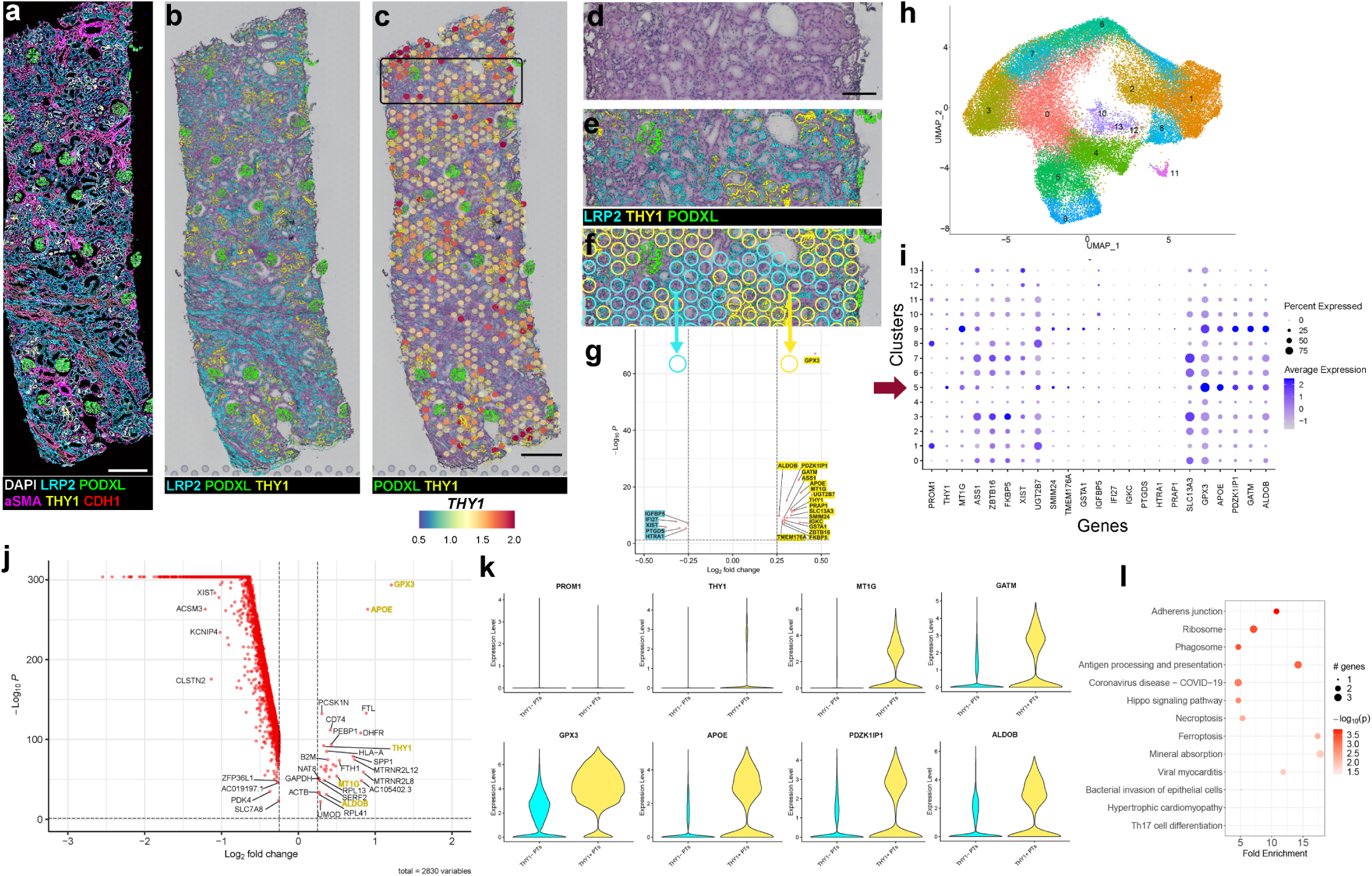
Bridging THY1 expression at the cell level from protein to RNA. Three sequential sections, adjacent to sections processed by CODEX, were processed by Visiumv1 spatial transcriptomics. (a) Representative CODEX image of a reference tissue showing 7 markers. (b) LRP2, PODXL, and THY1 expression registered on one of the sequential ST section stained with H&E. THY1 co-labels a subset of PT (LRP2-positive). (c) THY1 gene expression by ST registered with protein expression in CODEX. Clear/blue spots = minimal gene expression. Red/orange = higher gene expression. The black rectangle corresponds to (d-f). (d) H&E image from ST. (e) CODEX with LRP2, PODXL, and THY1 registered upon the ST H&E image. (f) Yellow circles correspond to ST spots overlaying regions that are positive for LRP2 and THY1 proteins. Cyan circles correspond to ST spots overlaying regions that are LRP2-postive and THY1-negative protein. (g) Volcano plot of ST differentially expressed genes (DEGs) comparing the two sets of spots from combined ST sections defined in (f), (Yellow vs. Cyan, 3 sections total). (h) UMAP of PT class from KPMP snRNAseq kidney atlas of health and disease (KPMP snRNASeq Atlas) clustered based on highly variable gene expression. (i) gene expression profile of the PT clusters from (h) displaying the genes uncovered in ST spots in (g) and PROM1. Arrow indicates a subcluster with high expression of THY1. (j) Volcano plot comparing gene expression between THY1-positive PTs compared to all other PTs in the KPMP snRNASeq Atlas. (k) Violin plots comparing the expression of a subset of genes from (j) THY1-positive vs. THY1-negative PTs. (l) Pathway analysis of DEG in THY1-positive vs. THY1-negative PTs. Scale bars: 500 μm in a & c, 200 μm in d.

### Changes of THY1 and PROM1 expression in proximal tubules during kidney disease

We compared the changes in PT cell types identified by CODEX between reference and disease (Figure 4a-c). THY1 expressing PT cells were significantly decreased in disease (Figure 4d). To validate these findings and further increase the robustness of our results, we examined THY1 and PROM1 expression in PTs in a larger validation cohort (n= 49 separate specimens with 2.35 million cells total) using biopsies from the Kidney Precision Medicine Project (KPMP) and healthy nephrectomy tissue from the Human Biomolecular Atlas Program (HuBMAP) (Supplemental Table 3). These tissue specimens were imaged with the newer Phenocycler system. Similar analysis was performed to uncover subtypes of PTs (n= 441002 cells) from the total cell clusters (Figure 4e-h and Supplemental Figure S4-S6). Because of the larger sample size of this validation cohort, disease biopsies were separated by diagnosis: reference, AKI or CKD. Similar to the CODEX discovery cohort, we observed that THY1-positive PTs were highly abundant in reference tissue, comprising ∼40 % of PT-cells, but they decreased significantly in AKI and CKD (Figure 4h-n). Remarkably, in healthy reference tissue sections, the expression of THY1 was consistently absent in S3 PTs in the medullary rays (Figure 4i & 4l), suggesting that the expression of THY1 at baseline is primarily localized to S1 and S2 tubules. Unlike THY1, the proportion of PROM1 expressing PT cells was trending higher in the CODEX data (Figure 4d) and was significantly increased in the Phenocycler data in AKI but not CKD (Figure 4h, j, k, m & n).

**Figure 4:**
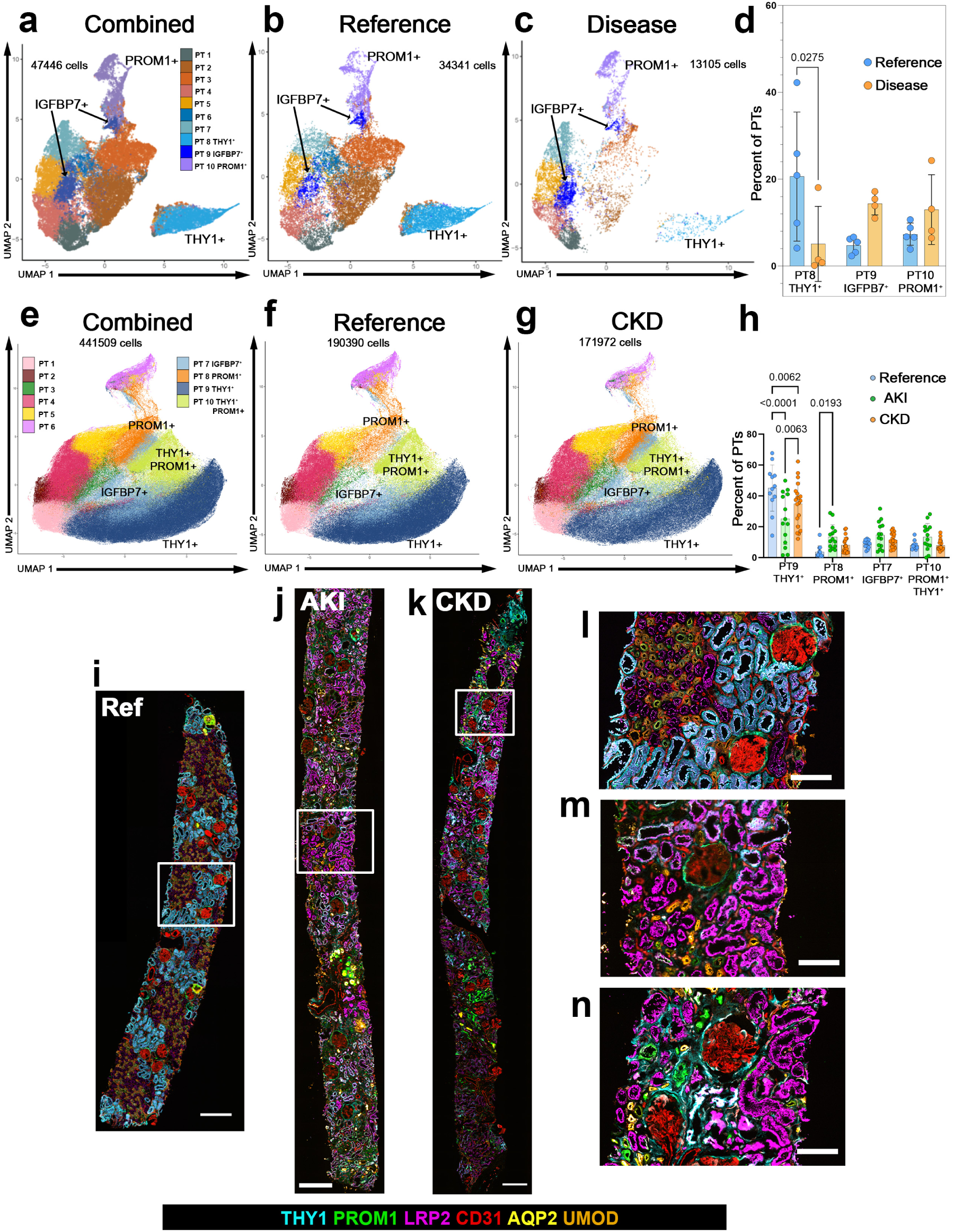
Changes in THY1 expression in disease. (a) Integrated UMAP of PT cells from combined CODEX dataset, separated into reference or disease groups (b and c, respectively). (d) Quantitative analysis of cell proportions in reference and disease tissue sections. (e-g) Integrated UMAPs of PT cells from combined Phenocycler dataset visualized based on condition: Combined, Reference, or CKD. Quantitative analysis of cell proportions in reference and disease conditions. (i-k) representative large-scale images of the Phenocycler data showing THY1, PROM1 and markers for endothelium (CD31), PTs (LRP2), Collecting ducts (AQP2) and TAL (UMOD); scale bars = 500 μm. (l-n) Insets of the boxed areas from (i-k); scale bars = 200 μm.

### CODEX-based cell trajectory analysis relative to THY1 and PROM1 expression

To understand whether the THY1 and PROM1 states of the PT represent steps along a spectrum or unique divergent cell processes, we next studied cell trajectories of proximal tubules in the CODEX data which highlighted the dynamics of THY1 and PROM1 expression (Figure 5). The trajectories of proximal tubules (high LRP2) in reference specimens show distinct paths towards THY1 and PROM1 expression (Figure 5a-c, trajectories t1 & t2 for THY1, t3 & t5 for PROM1). This is consistent with distinct expression profiles (such as injury markers) of these 2 sub-populations outlined in Figure S3. Interestingly, some PROM1-positive PTs in reference tissue had a low level of THY1 expression (Figure 5b-c, and Figure S7). This low level of THY1 is transient and lost with increasing PROM1 expression along trajectory 3 (t3 in Figure 5d-e). In disease specimens, the expression of THY1 is markedly reduced (Figure 5f-g). The trajectories towards THY1 and PROM1 expression (trajectories t4 & t1) remain distinct. Further, a population of PROM1-positive cells with THY1 expression were absent in disease (Figure 5b-c versus Figure 5g-h).

**Figure 5:**
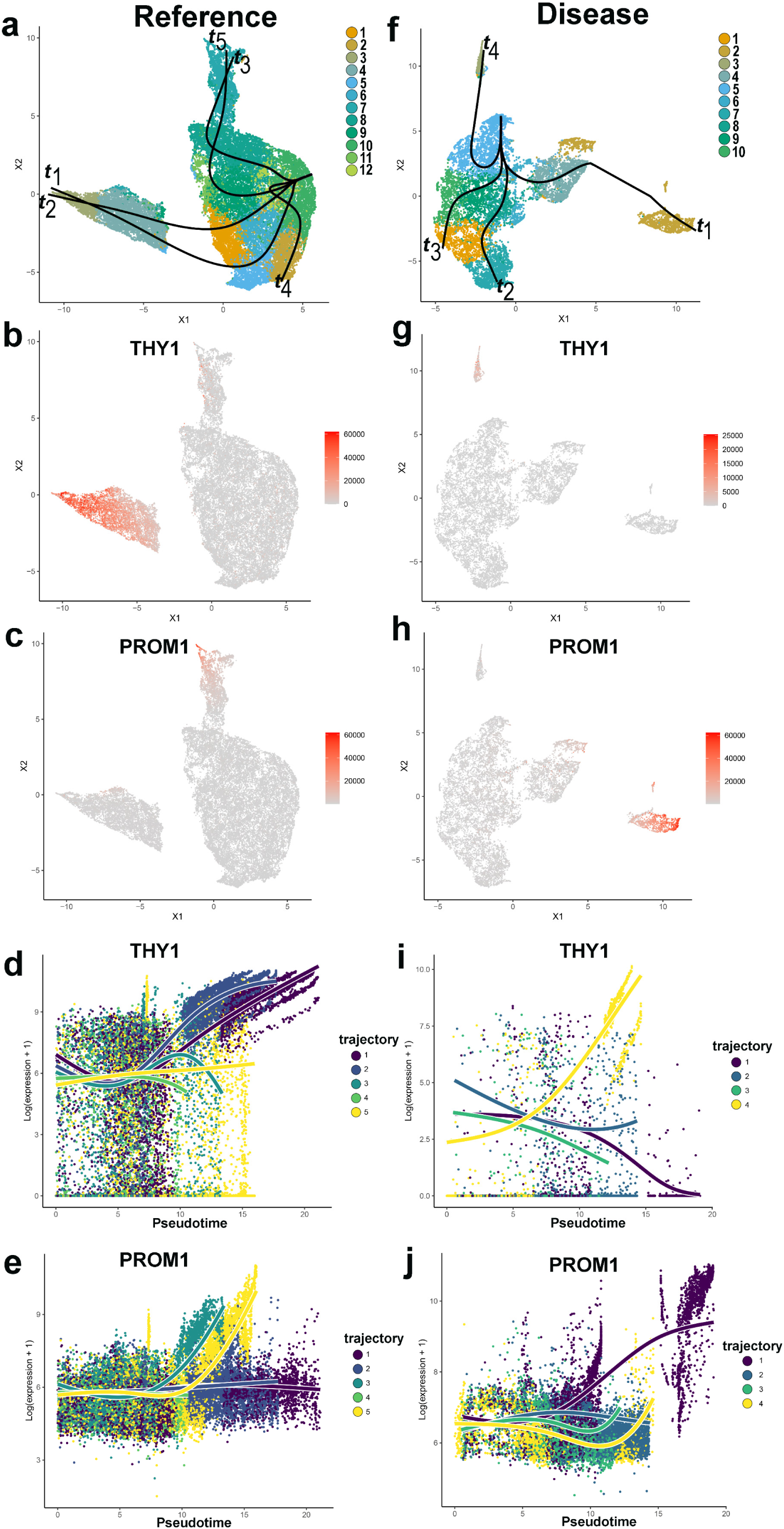
Trajectory analysis of PT cells accounts for the dynamics of THY1 and PROM1 expression in CODEX data. Reference (a-e) or disease (f-j) samples were separated and re-clustered and projected into UMAP space. (a and b) PT cell clusters from CODEX data in reference tissue with trajectory analysis (t1-t5) or in disease tissue (t1-t4), starting from clusters of PT with high LRP2 expression. (b & c or g & h) Feature plots for THY1 and PROM1 expression in reference and disease, respectively. (d & e or I & j) Pseudo-time spectrum of PT cells based on THY1 and PROM1 expression, in disease and reference samples respectively.

### snRNAseq-based cell trajectory analysis relative to *THY1* and *PROM1* expression

To determine if similar cellular trajectories are found in transcriptomics data, we studied the dynamics of the various populations of proximal tubular cells in the snRNA-seq dataset from the KPMP atlas organized by clinical cohort: reference, AKI, and CKD (Figure 6 and Figure S8). In reference tissue, there was a distinct population of *THY1*-positive PTs (cluster 3 in Figure 6a and Figure S8a) and 3 subpopulations of *PROM1*-positive PTs grouped together (cluster 4, 6 & 8). *PROM1*-positive subpopulations express varying degrees of injury markers (consistent with CODEX data Figure S3) such as *HAVCR1* (KIM1) and *VCAM1* (clusters 6 > 8> 4, Figure S8a). Notably, trajectories from healthy tubules towards *THY1*-positive and *PROM1*-positive PT cells are also distinct, like our CODEX data in reference tissue (Figure 5). In further concordance with our CODEX data, we observed a low level of expression of *THY1* in the *PROM1*-positive PT subpopulation (cluster 4) that expresses the least amount of injury markers (Figure 6a & Figure S8a, trajectory t3 Figure 6d & 6e).

**Figure 6:**
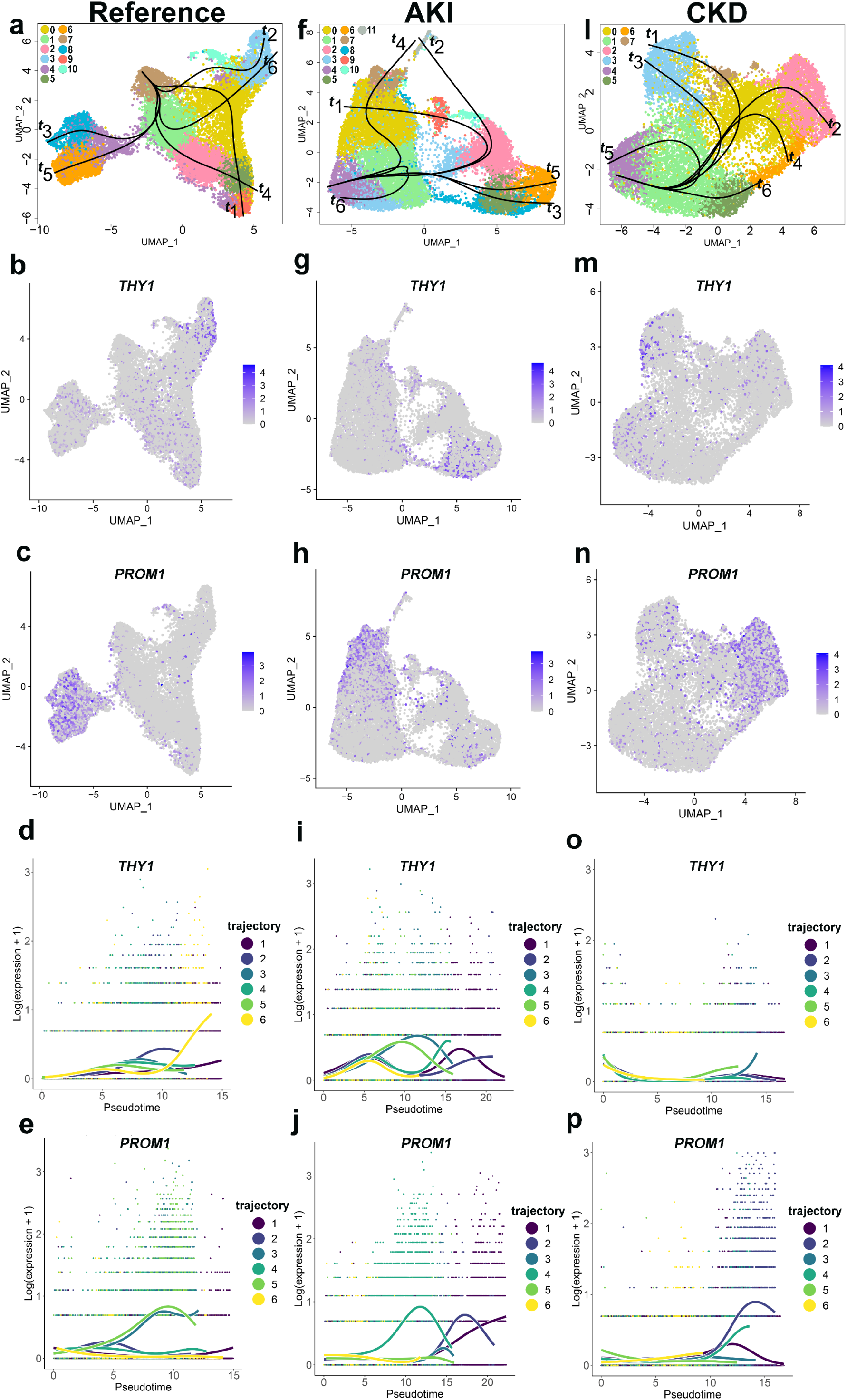
Trajectory analysis of PT cells accounting for the dynamics of *THY1* and *PROM1* expression in snRNAseq data. (a) PT cell clusters from the KPMP snRNAseq data in reference tissues with trajectory (t) analysis starting with cluster of PTs without any injury markers and expressing genes know to be present in healthy differentiated PT cells. (b & c) Feature plots for THY1 and PROM1 expression in reference PT cells. (d-e) Pseudo-time spectrum of PT cells based on THY1 and PROM1 expression, respectively, for each of the trajectories shown in (a). (f-j) the same analysis performed in (a-e) was done on cells from AKI tissue specimens. (k-p) Similar analysis performed on cells from CKD tissue specimens.

In AKI, the main population of *THY1* expressing PTs (cluster 5, Figure 6f & g and Figure S8b) remains distinct from *PROM1* expressing PTs (clusters 0 and 7, Figure 6f & 6h and Figure S8b), with separate trajectories from healthy PTs towards those with high *THY1* or *PROM1* expression (trajectories t3 & t5 vs. trajectories t1, t2 & t4, respectively, Figure 6f, i & j). High *THY1* PTs were characterized by near absent *HAVCR1* expression, but high *VCAM1*. *HAVCR1* was closely linked to *PROM1* expressing cells (clusters 0 and 7). There were also several subpopulations of PTs with intermediate levels of *PROM1* and *THY1* expression (Figure S8b & d), particularly on trajectories t1, t2 and t4 (Figure 6i-j) from healthy PTs to high *PROM1* expressing cells. This suggests an expression spectrum leading to an injury state characterized by low *THY1* and high *PROM1* and *HAVCR1* expression.

In CKD, the expression of *THY1* is markedly reduced and only observed at low levels in 2 subpopulations (clusters 3 & 4, Figure 6l & 6m and Figure S8c) that have reduced *HAVCR1* or *VCAM1* expression. Conversely, the expression of *PROM1* remained significant and clearly distinct from *THY1* expression (clusters 2 & 6, Figure 6l & 6n and Figure S8c), with distinct trajectories from healthy towards PTs with high *THY1* or *PROM1* expression (trajectory t3 vs. trajectories t2 & t4, respectively, Figure 6l, o & p). PT expressing *PROM1* also had high *HAVCR1* and *VCAM1* expression, suggesting a shift towards a maladaptive repair phenotype^11,13,16^.

Cumulatively, the cell expression and trajectory data from CODEX and snRNAseq are concordant and suggest that THY1 and PROM1 are likely on an opposite spectrum of disease progression with overlapping intermediary states during acute injury. Specifically, the loss of THY1 and shift towards high PROM1 states are associated with CKD.

### Neighborhood analysis from the CODEX imaging data

To determine the cellular microenvironments for the distinct THY1 and PROM1 PT subpopulations, we performed a cell-centric neighborhood analysis^7,27^ which uncovered 19 cell neighborhood clusters or niches where specific types of cells significantly associated in the same neighborhoods (Figure 7). When considering the predominant cell types present within each niche and their relationships to other niches in a dimensionally reduced space (Figure 7a-b) we deconstructed the niche landscape into nephronal, immune, endothelial and stromal microenvironments. For example, we can specify niches enriched in glomerular cells (N15 > N5, likely glomerular or periglomerular environments), PTs (N2, N3, N4, N6, & N8), immune cells (myeloid: N5>N18>N7>N17; T cells: N18>N7>N17) and fibroblasts (N7). THY1-positive PTs and PROM1-positive PTs account for the largest proportion of cells in the N3 and N17 niches, respectively. Notably, the abundance of immune cells in N3 is in the lowest tertile compared to N17 niche which is in the highest tertile of immune cell abundance within all the niches (Figure 7b), suggesting an association between PROM1-positive PT and immune cell activity. Comparing the distribution of niches in disease vs. reference, we observed an increase in fibroblast (N7) immune (N5, N18) and PROM1-positive PT (N17) niches and a decrease in the THY1-positive PT (N3) niche (Figure 7c). Pairwise analysis further confirmed the positive association of PROM1-positive PT and negative association of THY1-positive PTs with immune cells (Figure 7d). These data are consistent with the observed changes in PT cell composition and trajectories in health and disease detected in both the CODEX and snRNA-seq analysis.

**Figure 7:**
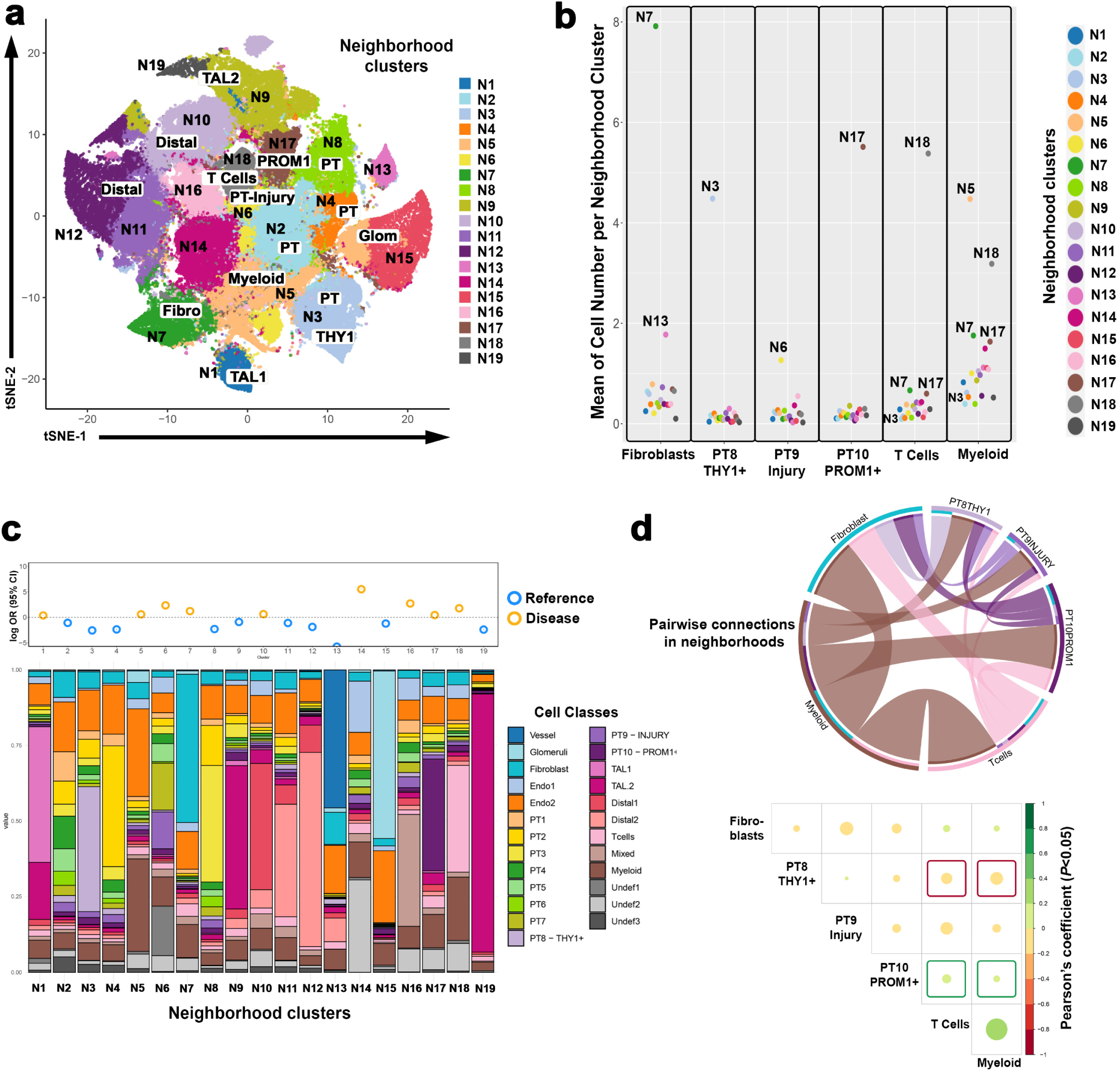
Neighborhood analysis in CODEX data highlights epithelial immune cell interactions. (a) tSNE plot showing neighborhood clusters or niches (N1-19, each dot is a niche) based on the average distribution of cell types in each niche. The major cell type or tubular type in underlying niches is indicated. (b)Distribution for specific cell types in all neighborhood clusters, to identify the niches with the largest numbers of specific cell types. (c-top) Distribution of cell types in neighborhoods and the propensity of niches to be found in reference or disease as assessed by odds ratio and (c-bottom) the average distribution of cell types in each niche. (d) Interactions of PT, immune cells, and fibroblasts in niches, top: chord plot to visualize the pairwise cell-cell interactions in all niches, bottom: pairwise correlation between cell types in all neighborhoods by Pearson’s coefficient. Red boxes highlight negative correlation of THY1-positive PTs with immune cells, and green boxes highlight the positive correlation between the immune cells and PROM1-positive PTs.

### Cell-cell interaction analysis

To further characterize the functional significance of *THY1* expression in PTs, we performed ligand-receptor analysis between subtypes of PTs and all cells in the KPMP snRNA-seq atlas (Figure 8). Our results demonstrate that some PT populations are prone to interact with epithelial, endothelial and immune cells using basement membrane ligands such as collagen IV subunits that interact with various integrins (Figure 8a-c). Such interactions are key for attachment to extracellular matrix components, cell growth and differentiation ^28,29^. These ligand receptor interactions are most pronounced for *PROM1*-positive PTs, which also have other interactions that are consistent with potential immune activation (CD226 activation^30^) and angiogenesis (THBS1 signaling^31^), particularly in disease. Conversely, these interactions are markedly absent in *THY1*-positive PT cells (Figure 8a-d). *THY1*-positive PTs only retain SPP1-linked cell interactions, which may be^32^. Cumulatively, these findings suggest a potential role of *THY1*-positive PTs in de-differentiation and transmigration, which are key features of cells that participate in regeneration.

**Figure 8:**
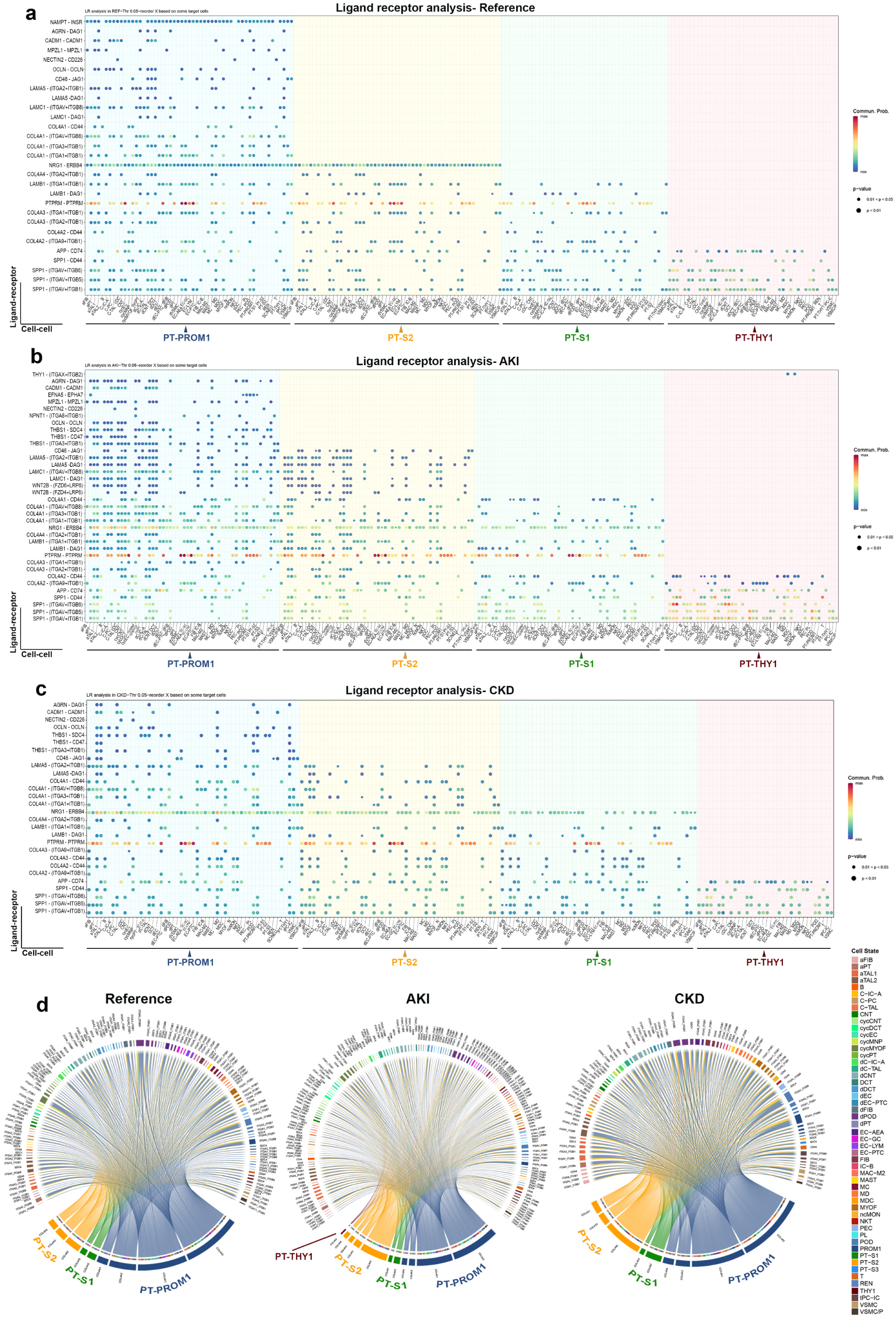
Ligand receptor analysis for PT cells from snRNAseq data. (a-c): Bubble plots from reference, AKI and CKD tissues, respectively, showing the ligand receptor interactions for PT cell subtypes on the y axis, and specific cell-cell interactions on the x axis. (d) chord plots highlighting collagen 4 subunit interactions in the various subtypes of PT cells.

## Discussion

In this work, we characterized subpopulations of proximal tubules (PTs) in the human kidney expressing THY1 and PROM1, markers linked with repair and regeneration. We localized these subpopulations with spatial gene and protein expression and further identified a *THY1*-positive PT subcluster within a snRNA-seq atlas, characterized by regenerative pathways and distinct from the PROM1-positive PTs. We then quantified changes in THY1-positive and PROM1-positive PT cells and charted their trajectories in health and disease. Broadly, THY1 and PROM1 are considered markers of stem cells and are thought to modulate tissue response to injury^19,21^. Whereas persistent expression of PROM1 in renal epithelial cells has been associated with maladaptive repair and progression towards fibrosis^1^, the role of THY1 and changes in its expression in PTs is less well understood.

In non-epithelial cells such as fibroblasts, leukocytes, neurons and endothelial cells, THY1 has been implicated in modulating processes involved in tissue regeneration such as cell migration, proliferation and matrix remodeling^21,22,33-35^. It is known that THY1 is not usually expressed in murine kidney epithelium^24^, and this has been further confirmed by the publicly available single cell transcriptomic databases ^36^. Therefore, the low expression reported in some mouse models^37^ likely originates from other cell types such as immune, endothelial or stromal cells. Interestingly, induced expression of human THY1 in murine kidney causes proliferative abnormalities of proximal tubules^24^, thereby supporting the posited role for THY1 in tissue regeneration. Contrary to the mouse, and consistent with this work, Wu et al recently reported constitutive THY1 protein and RNA expression in the healthy human kidney tubulointerstitium, which was significantly decreased in diabetic kidneys^23^. Single cell RNAseq databases of human kidney confirmed the expression of THY1 in PTs along with other cells such as fibroblasts and immune cells^23^. Surprisingly, increased levels of soluble urinary THY1 were also associated with kidney disease, which was attributed to possible increased shedding of THY1.

Our data conclusively demonstrates that in reference kidney tissue, approximately 20-50% of PTs (based on the imaging platform used) express THY1. For unclear reasons, the ubiquity of this cell state was underrepresented in the snRNA-seq atlas and underscores the important complementary roles of spatial protein imaging and transcriptomics. A much smaller proportion of PTs express PROM1 in healthy states. Further, THY1 loss in PTs and shift towards states of high PROM1 expression is clearly associated with kidney disease. THY1-positive and PROM1-positive PTs are likely on opposite ends of a spectrum of disease progression with some overlapping intermediary states during acute injury. Our molecular cell-cell interaction analysis is consistent with a regenerative phenotype in THY1-positive PT cells, and pro-inflammatory features for PROM1-positive PT cells. This was further supported in the neighborhood analysis where niches enriched with PROM1-positive PTs had high abundance of immune cells. Our data support a model in which THY1 loss and the shift towards PROM1 expression in PTs could be a valuable indicator of kidney disease progression. Therefore, we propose that dual THY1 and PROM1 staining in kidney biopsies could serve as a staging tool for chronicity and a prognostic biomarker for kidney disease progression. This will require further validation studies in larger cohorts.

There are several novel approaches and findings in this work that are worth highlighting. In a robust pipeline, we integrated large scale CODEX multiplexed imaging data from multiple specimens in a single analytical space. Although a spatial protein imaging-based analysis was recently described in diabetic kidneys^38^, our approach benefited from cross-validation with orthogonal techniques and datasets. We identified unique PT clusters that express either THY1 or PROM1 with high confidence, amidst other PT subtypes that were classified based on their protein expression profile. In additional analysis, PROM1-positive PTs also expressed other markers of injury, consistent with transcriptomics data showing that *PROM1*-positive PTs express markers of failed repair, especially in CKD. THY1 and PROM1 PTs had reciprocal patterns of expression in reference and disease, except for intermediary states that were seen in reference and AKI.

To increase the robustness and generalizability of our findings in the CODEX dataset, we used a large Phenocycler (newer iteration of CODEX) validation cohort. In this larger cohort we validated the findings from the initial CODEX discovery cohort, showing a significant decrease in THY1-positive PTs and an increase in PROM1-positive PT population in disease.

A strong agreement and alignment between protein and RNA expression data was demonstrated by the integrated analysis based on co-registration of CODEX and ST on consecutive tissue sections. Not only did this analysis confirm the concordance of THY1 protein and RNA expression in PTs, but it also allowed us to investigate protein-defined THY1-positive PTs at full transcriptome depth. This integrative analysis could be adopted for other molecules or even other tissues when consecutive sections are available for interrogation using spatial protein imaging and spatial RNA expression. The concordance of the ST-derived gene expression profile of THY1-positive PT cells in the CODEX data with the transcriptomics data from the KPMP atlas, which were generated independently, increases the robustness of the findings.

This work also provides an innovative analysis charting the trajectories of PTs based on spatial protein imaging. These trajectory and pseudo time analysis in CODEX and snRNA-seq yielded similar results. Such congruent findings by independent technologies from 2 separate datasets not only enhance the robustness of the results, but also support the usefulness of trajectory analysis in the CODEX data, especially when validated by snRNA-seq, which is considered a gold standard for such analyses^39^.

Integrated neighborhood analysis on the CODEX data is also novel and showed a negative association between THY1 and immune cells, whereas PROM1-positive PTs were enriched within neighborhoods that have significant immune activity. These findings could be consistent with a failed repair phenotype of PROM1-positive PTs, which are associated with immune activation and expression of injury markers such as KIM1 and VCAM1^11-14,16^. The cell chat analysis using the snRNA-seq data was also consistent with these observations and further extended our understanding of potential interactions of *THY1*-positive and *PROM1*-positive PTs with other cell types. *THY1*-positive PT cells have remarkably reduced receptor ligand interactions, which were mostly centered around interactions with SPP1 (Osteopontin). Such interactions have been linked to dedifferentiation and transmigration^32^, which could be part of the regenerative profile associated with THY1. Conversely, *PROM1*-positive PTs exhibited extensive ligand receptor interactions with epithelial, endothelial and immune cells using collagen IV subunits that could engage various integrins. These findings support a pro-injury and inflammatory profile of PROM1-positive cells. The consistency of findings from the snRNA seq data cell-cell interaction with the CODEX neighborhood and trajectory analyses (from CODEX and snRNA seq) is quite remarkable and provide cross validation across various assay platforms and datasets.

Our studies have a few limitations that are worth discussing. The CODEX discovery cohort has a relatively small sample size, which could be limiting in terms of sampling bias and generalizability. We mitigated this limitation by including a larger validation cohort, which reproduced and further extended the results. This larger validation cohort is currently being further analyzed and will be the subject of future work. Furthermore, the concordance of the CODEX data with the transcriptomics data validates the findings and the robustness of the results. Based on our data we cannot infer causality on the role of THY1 or PROM1 PTs in repair or regeneration, but the existing experimental data from other groups for both THY1 and PROM1 strongly support our conclusions^19,24^.

In summary, we show for the first time that protein imaging modalities such as CODEX offer a robust platform for data analysis that could complement existing RNA-based platforms. Our data point to THY1 and PROM1 as important players in proximal tubule biology and could serve as useful biomarkers for disease staging and progression as well as potential therapeutic targets.

## Methods

### Tissue sources and study approvals

Ethical compliance: We have complied with all ethical regulations related to this study. All experiments on human samples followed all relevant guidelines and regulations. The relevant oversight information is given based on tissue sources (detailed below).

#### CODEX discovery cohort

Detailed analyses were done using CODEX imaging on a cohort of biopsies or reference kidney tissue consisting predominantly of cortex. Human kidney tissue biopsies and nephrectomies were preserved in Optimal Cutting Temperature (OCT) medium and were acquired from the Biopsy Biobank Cohort of Indiana, under a waiver of informed consent as approved by the Indiana University IRB (IRB #1906572234). Reference nephrectomy tissues were also obtained from the Kidney Precision Medicine Project central biorepository. Three reference kidney tissues, also preserved in OCT, were obtained via percutaneous nephrolithotomy carried out in patients with stone disease, as part of an ongoing study (Indiana University IRB Protocol #1010002261). For relevant demographic information about each tissue donor, the corresponding tissue source and data availability, please see Supplemental Table 1.

#### Phenocycler validation cohort

Kidney biopsies imaged with Akoya Biosciences Phenocycler-Fusion 2.0 from the KPMP atlas were used for validation of the changes of THY1 and PROM1 staining in proximal tubules. Imaging data are publicly available in the KPMP atlas. Human samples collected as part of the KPMP consortium were obtained with informed consent and approved under a protocol by the KPMP single IRB of the University of Washington Institutional Review Board (IRB#20190213). We also used reference tissue from the Human Biomolecular Atlas Program (HubMap) (https://hubmapconsortium.org/hubmap-data/). Samples as part of the HuBMAP consortium were collected using informed consent by the Kidney Translational Research Center (KTRC) under a protocol approved by the Washington University Institutional Review Board (IRB #201102312). Detailed analysis of this cohort will be published elsewhere as part of a Spatial Atlas output from KPMP.

#### Single nuclear RNA sequencing sample source

A combined snRNAseq atlas from the Human BioMolecular Atlas Program (HubMap) (https://hubmapconsortium.org/hubmap-data/) and Kidney Precision Medicine Project (KPMP) (https://www.kpmp.org) datasets (200,338 nuclei) with 41 samples from 36 subjects was used^1^. This dataset was previously published and publicly available (GSE183277).

#### Spatial transcriptomics sample source

2 Reference tumor nephrectomy tissues were also obtained from the Kidney Precision Medicine Project central biorepository (Supplemental Table 1).

### Antibody Selection, Conjugation and Validation

#### CODEX

Following a careful selection of structural, immune and injury markers, based on available literature, public data domains, and previous work, we composed an initial panel of 23 markers that was applied to all specimens. This panel was subsequently extended to include more specialized markers (total 38) to label cell types, and cell states (see Supplemental Table 1 for marker panel used for each specimen and Supplemental Table 2 for details about the panels).

Twenty-six of the antibodies from the final panel were conjugated in-house using the protocol outlined by Akoya Biosciences and Black et al^40^. Commercially purchased antibodies first underwent a reduction step using a Reduction Master Mix (Akoya Biosciences). Lyophilized barcodes were then resuspended using Molecular Biology Grade Water and Conjugation Solution. The barcode solution was then added to the reduced antibody solution and incubated for 2 hours at room temperature. After incubation, the newly conjugated antibody-barcode was purified in a 3-step wash/spin process and stored at 4*C. Successful conjugation was validated via gel electrophoresis as well as immunofluorescent staining and confocal imaging.

#### Phenocycler

The antibody panel used in CODEX was further expanded to 42 markers. 41 markers used in this analysis (Nestin was excluded) have been validated and published as part of Organ Mapping Antibody Panels (OMAP)^41^ (https://doi.org/10.48539/HBM542.NNZP.924) and shown in Supplemental Table 4.

### Tissue Preparation and Imaging

#### CODEX

Ten µm thick human kidney tissue sections were cut from OCT blocks onto poly-L-lysine coated square coverslips, which were subsequently processed using the protocol outlined by Akoya Biosciences and as described previously by Golstev et al^42^ and Melo Ferreira et al^8^. Antigen retrieval was conducted with a 3-step hydration process, followed by fixation with 1.6% PFA after initial fixation, an antibody cocktail of the markers listed in Supplemental Table 2 was dispensed among the coverslips. The antibody solution was left on the coverslips overnight at 4°C. The following day, the staining solution was washed from the tissues, and a second fixation was performed. Oligonucleotide probe staining and fluid handling were performed with the CODEX system from Akoya Biosciences. Automated tile-scan imaging of the tissue between probe staining rounds was performed on a Keyence BZ-X810 microscope fitted with a 20X objective. The resulting images were processed using the CODEX Processor (Akoya Biosciences) and visualized using FIJI/ImageJ^43^.

#### Phenocycler

Tissue processing for the Phenocycler follows the same protocol as described above for the CODEX system, but the poly-L-lysine coated coverslips are replaced by SuperFrost Gold+ Charged slides. After staining slides were covered with an Akoya Biosciences FlowCell. Cover-slipped slides are incubated in 1X CODEX Buffer with Buffer Additive (AKOYA) for 10 minutes to ensure proper adhesion of the FlowCell to the slide. Slides were either kept in storage buffer until ready for imaging or imaged immediately after the incubation. Imaging of the tissues was conducted with a 20x objective fitted on the Akoya Biosciences Phenocycler 2.0 microscope and fluidics handler. Image stitching and processing was also performed with the Phenocycler 2.0.

### Image Analysis

#### CODEX

CODEX images were brought into ImageJ/FIJI and analyzed using a custom-made plug-in: Volumetric Tissue Cytometry and Exploration (VTEA)^7,44^. Nuclei stained with DAPI were segmented with VTEA. A morphological dilation around each segmented nuclei was used as a proxy segmentation of cytosol. Using these nuclear or cytosol segmentations, the mean intensity in aligned images from additional CODEX rounds were calculated to detect protein markers in the putative nuclei and cytosol of individual cells. The quality of individual protein markers was assessed on a channel-by-channel basis. Channels were excluded from the dataset with aberrations including, 1) no detectable signal, 2) signal correlated with background images, or 3) imaging artifacts (e.g. exposure inconsistence across image tiles). Further, the semi-unsupervised clustering of segmented cells with measured protein markers were used to help identify aberrations (e.g. cell clusters that correlated with inconsistent exposures across tile scanning). For this, cells were projected into *t-*SNE space with a learning rate of 100 and a perplexity of 40, and clustering was conducted using Ward Hierarchal Clusters based on the same feature space, with a maximum of 15 clusters and mapped back to the image to identify correlations with artifacts.

After tissues were processed individually and artifacts removed, measurements from segmented cells were imported into R and combined into a single dataset following Z-scaling. Combined datasets were clustered using the Louvain methods and a feature space including the mean intensity of CD31, CD3, CD45R0, CD4, CD45, AQP1, CD11c, THY1, a-SMA, E-Cadherin, PROM1, Podocalyxin, LRP2, CD68, UMOD, IGFBP7, and Cytokeratin 8. After labeling the major clusters, the proximal tubule clusters were extracted as subsets and re-labeled using a tubular epithelium specific set of features that included the mean intensities of AQP1, THY1, a-SMA, E-Cadherin, PROM1, LRP2, CD68, UMOD, IGFBP7, Cytokeratin 8.

#### Phenocycler

Final “qptiffs” images generated by Phenocycler-Fusion (AKOYA) were imported into ImageJ/FIJI for cropping biopsies into individual image files. Segmentation of nuclei based on DAPI staining was done using VTEA, as with the CODEX discovery cohort. Cortical and medullary regions were defined for each tissue to be used as additional features in downstream analysis. The resulting segmentation from all biopsies was imported into R, and data were z-scaled. Forty-one markers and the region feature (cortex or medulla) were used to cluster the scaled data using FastPG (https://github.com/sararselitsky/FastPG), with k = 100. After clustering, UMAP embeddings were calculated, and data plotted along with violin plots of both scaled data and raw intensity values to identify clusters. Cluster identities were further validated by mapping back the clusters onto representative tissue sections. Consistent with the CODEX analytical pipeline, PT clusters (based on LRP2 and AQP1 expression) were extracted into subsets and re-clustered. The feature space for this subclustering was the same as the one used in the CODEX discovery cohort in addition to the region (cortex/medulla). Clustering was done using FastPG with a k = 500, which still yielded a high number of clusters because of the larger sample size. Redundant clusters based on the expression of THY1, PROM1 and IGFBP7 were further grouped together.

### Cell neighborhood analysis in CODEX data

The proximal tubule subclusters were imported back into VTEA for neighborhood generation and analysis^7^. The “Spatial by cell” method in VTEA was used, with a neighborhood diameter of 55 µm to match the spot size used in VISIUM spatial transcriptomics. VTEA generates the counts (“Class Sums”) and the fraction-of-total of each cell-type for each neighborhood which were imported into R for further analysis. To determine which cell types were spatially associated in pairs, correlation matrices were calculated for all samples (corrplot v. 0.92). Chord plots (circlize v. 0.4.16) were also generated for cell types of interest, informed by the correlation matrices. For neighborhood analysis, the “Class Sums” were clustered with Louvain (igraph v. 1.5.1) and *t*-SNE (Rtsne v. 0.16) embedding and neighborhood cluster or niches identified. Stacked bar plots were used to visualize the distribution of cell types within each niche, and the odds ratios were calculated to assess the prevalence of a given niche in the reference versus disease cohorts.

### Spatial Transcriptomics (ST)

ST was done on three 10 um sections obtained from 2 human reference kidney tissue specimens. The preparation and imaging of tissues were done using Visium Spatial Gene Expression protocol for polyA capture in fresh frozen tissue (10x Genomics, CG000240 protocol, Visium Tissue Preparation Guide) ^8^. Using a Keyence BZ-X810 microscope equipped with a Nikon 10x CFI Plan Fluor objective, hematoxylin and eosin (H&E) stained brightfield mosaics were collected and stitched prior to mRNA capture. mRNA was extracted from tissue after 12 minutes of tissue permeabilization. Isolated mRNA was captured by oligonucleotides in the fiducial capture areas and reverse transcribed. Libraries were prepared based on the Visium 1.0 protocol (10X genomics CG000239 protocol) which included, second-strand synthesis, denaturation, cDNA amplification, and cleanup using SPRIselect (Beckman Coulter). Multiplexed sequencing was performed on a NovaSeq6000 (Illiumina). Sequencing data was demultiplexed and mapped to the reference genome GRCh38 3.0.0, and counted in Space Ranger (10X Genomics, v1.0.0). Data processing was performed in Seurat (v.4.4)^45^.

snRNAseq kidney atlas data, generated by KPMP and HuBMAP consortia ^1^ were mapped to the ST. SCTransform was used for reads normalization^45,46^. Additionally, dimensionality reduction was done with Seurat’s implementation of uniform manifold approximation and projection (UMAP) and the expression signatures of spots for cell types were calculated and mapped using a transfer score system based on Seurat anchors method^45^. This score shows the association of the transcriptomics profile of each spot with a specific cell type. A higher transfer score implies that a greater proportion of a specific cell type is mapped to a particular ST spot.

The expression datasets, obtained from the mappings of each section, included the gene counts and sample metadata. Three datasets from each section were merged and a UMAP was regenerated. Batch correction was accomplished by normalizing the raw counts of the merged object with SCTransform ^45,46^.

### co-registration and integrated analysis with CODEX

To co-register ST and CODEX, 3 consecutive ST and CODEX sections from 2 separate reference tissue specimens were used. In the CODEX data, LRP2, THY1, and Podocalyxin (PODXL) staining were used to outline proximal tubules (PT), THY1-positive PTs, and glomeruli, respectively.

ST-CODEX alignment was done manually using photoshop (PS v.23.3.1). CODEX images composed of LRP2, THY1 and PODXL were aligned with the H&E images of ST slide using linear transformations to align landmark structures, such as glomeruli and large tubules. Accordingly, other structures were overlayed precisely. Using this alignment, the spots in ST samples were categorized based on expression of THY1 and LRP2 in CODEX. Using loupe browser (v.6), four groups of spots were defined as LRP2-positive THY1-negative, LRP2-positive THY1-positive, edge layer of tissues, and the rest of the cells. The last two groups were excluded from the analysis.

To find the differentially expressed genes (DEGs) in the two groups of LRP2-positive THY1-negative and LRP2-positive THY1-positive spots, “FindMarkers” Seurat function was used with the p-value of 0.05 and average log2 fold change >0.25.

### snRNAseq analysis of PTs from KPMP atlas

The cortical subset of PTs including 29 specimens was taken from the snRNA-seq atlas^1^. To compensate for batch effects, the Seurat anchors method was applied to integrate samples in this cortical PT subset. PTs were clustered using Louvain algorithm with a resolution of 0.5. Based on *THY1* differential expression, cluster 5 was selected as a THY1-positive subclass of PT cell. Using “FindMarkers” function, the differentially expressed genes were found in this cluster compared to other 13 clusters. Pathway enrichment was done for the top 33 upregulated and top 33 downregulated genes. The R package pathfindR ^47^ was used for pathway analysis using the KEGG database.

### Trajectory analysis

Trajectory and pseudotime analysis of CODEX and snRNASeq data were performed in R using either the slingshot^48^ (v.2.6.0) and tradeSeq^49^(v.1.12.0-1.18.0) packages. The pseudotime spectrum of PT cells from PT clusters with high LRP2 expression to *THY1*-positive and *PROM1*-positive was measured using tradeSeq package.

#### CODEX

Trajectory analysis was performed on CODEX PT sub-clusters from either reference or disease using the average expression of AQP1, THY1, alpha-smooth muscle actin, E-cadherin, PROM-1, LRP2, CD68, UMOD, IGFBP7, and Cytokeratin-8 starting with the cluster with the highest average LRP2 expression. Trajectory and pseudotime analysis were performed independently in reference and disease.

#### snRNAseq

PT cells were subsetted from the snRNASeq datasets. Reference, AKI and CKD cohorts we separated and independently integrated using the Seurat anchors method. PT subtypes within each cohort were identified with a clustering resolution of 0.5. Trajectory and pseudotime analysis was performed independently in reference, AKI, and CKD. PTs with high LRP2 and other PT differentiation markers and low injury markers were considered the starting point of trajectories.

### Cell-Cell Communication

We evaluated the interactions of cells in *THY1*-positive*, PROM1*-positive, S1 or S2 PTs with other kidney cell types in snRNAseq atlas using <u>CellChatDB</u> package (v 1.6.1) ^50^. This approach defines ligands on the source cells interacting with receptors on target cells.

### Statistics

Unless otherwise specified, values are reported as mean +/- standard deviations. Bar graph scatter plots were generated using GraphPad Prism (10.0.2). Identification of differentially expressed genes between groups was performed with negative binomial generalized linear model. Significant genes were assigned using and adjusted p-value of <0.05 with Bonferroni correction. A two-way ANOVA model was used to compare the different cell proportions in the CODEX and Phenocycler data. For cell communication analysis, significance was derived from one-sided permutation test (the CellChat package default). Association of disease conditions with neighborhoods in CODEX was determined using odds ratio.

### Source data availability statement

Sequencing data generated for this study or previously unpublished are available at Gene Expression Omnibus (GSEXXXXXX). All imaging data and analyses are available on Zenodo (doi: TBD). Additional imaging data can be found at https://atlas.kpmp.org/. Source data for all Figures is available on Zenodo (doi: TBD). All code used to generate analyses and Figures available on Zenodo (doi: TBD).

## Funding/Acknowledgement

We would like to acknowledge the NIH common fund Human Biomolecular Atlas Project (HuBMAP) and the NIDDK’s Kidney Precision Medicine Project. This project was supported by an award from the Ralph W. and Grace M. Showalter Research Trust and the Indiana University School of Medicine. The KPMP is funded by the following grants from the NIDDK: U01DK133081, U01DK133091, U01DK133092, U01DK133093, U01DK133095, U01DK133097, U01DK114866, U01DK114908, U01DK133090, U01DK133113, U01DK133766, U01DK133768, U01DK114907, U01DK114920, U01DK114923, U01DK114933, U24DK114886. The HuBMAP work in this manuscript is supported by common fund grants U54DK134301 and OT2OD033753. Additional support for this work from 5U54DK137328.

## Author Contributions

Coordination of manuscript writing and project: M.A., D.B., A.R.S, P.C.D., S.W., T.M.E. Contribution to Patient Recruitment and Tissue Collection: S.J., K.J.K, J.V., J.O., P.P., S.E.R., S.S.W., K.K.K., C.P., I.D., J.H., M.K. J.H. Contribution to Tissue Processing: W.S.B., M.F, J.H., Y-H.C, C.P. Contribution to Imaging data generation: W.S.B., A.R.S., D.B.,M.F., T.M.E., Contribution to ST data generation: R.M.F., Y-H.C., M.T.E., Contribution to image data analysis: M.A., A.R.S., D.B., P.S. S.W., P.C.D., T.M.E., Contribution to transcriptomics analysis: M.A, R.M.F., M.T.E.,S.J. M.K. S.W. P.C.D. T.M.E., contribution to data interpretation: all authors, Contribution to writing manuscript: all authors.

