## Supplemental Figures for "Integration of spatial multiplexed protein imaging and transcriptomics in the human kidney tracks the regenerative potential timeline of proximal tubules"

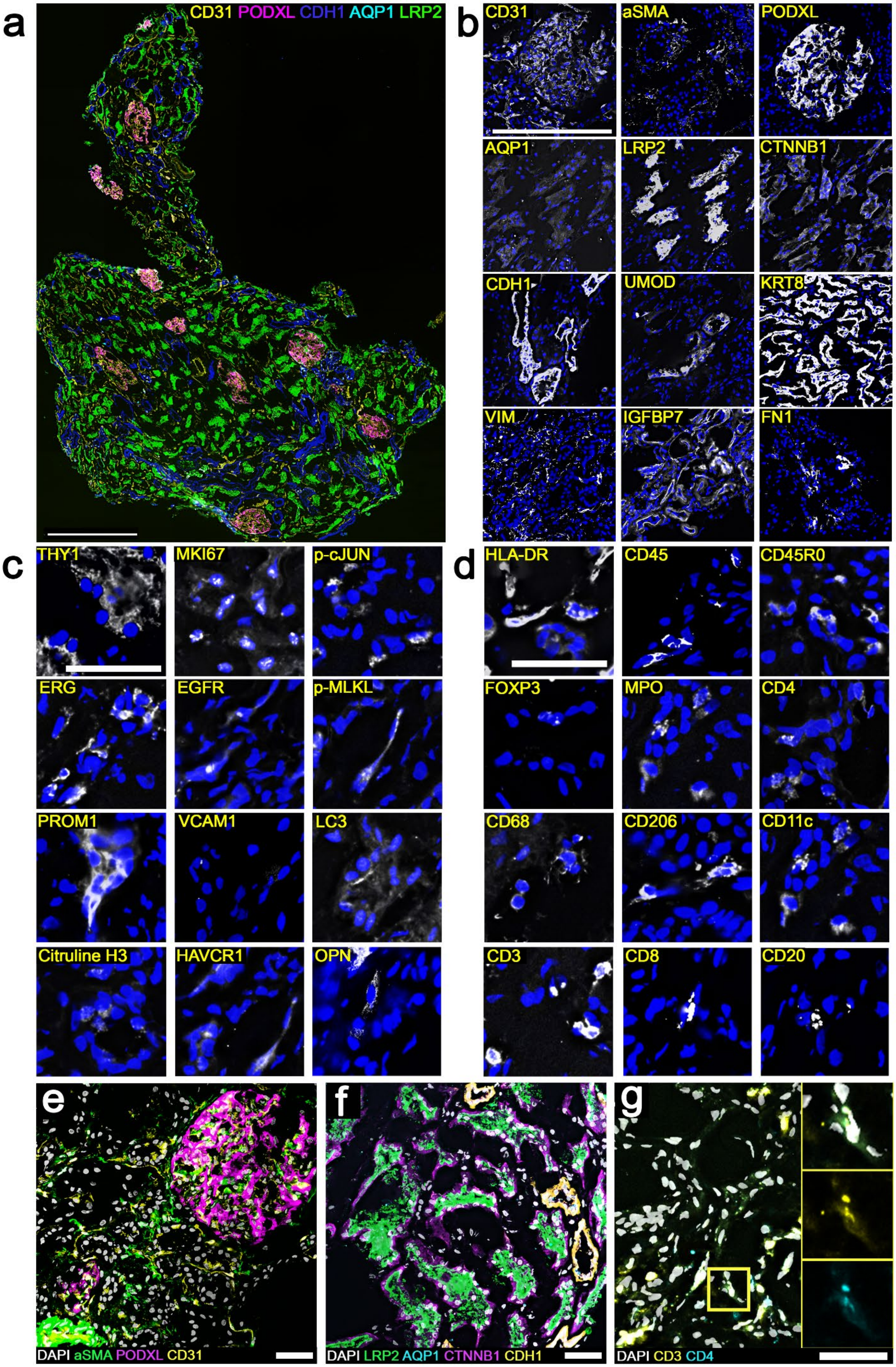
a
CD31 PODXL CDH1 AQP1 LRP2
b
CD31
aSMA
PODXL
AQP1
LRP2
CTNNB1
CDH1
UMOD
KRT8
VIM
IGFBP7
FN1
c
THY1
MKI67
p-cJUN
ERG
EGFR
p-MLKL
PROM1
VCAM1
LC3
Citruline H3
HAVCR1
OPN
d
HLA-DR
CD45
CD45R0
FOXP3
MPO
CD4
CD68
CD206
CD11c
CD3
CD8
CD20
e
DAPI aSMA PODXL CD31
f
DAPI LRP2 AQP1 CTNNB1 CDH1
g
DAPI CD3 CD4

**Supplemental Figure 1. CODEX multiplex imaging and validation of antibody panel**

a. CODEX imaging of structural markers [CD31 (vasculature), PODXL (podocytes, glomeruli), CDH1 (tubular epithelium), AQP1 (proximal tubules) and LRP2 (proximal tubules)] in a reference kidney tissue. Scale bar is 500 μm. b-d. Staining patterns or individual structural (b, scale bar =500 μm), cell states (c, scale bar = 200 μm) and immune markers (d, scale bar = 200 μm). e. Composite image of endothelial and stromal stains. Scale bar = 50 μm. f. Composite image of both proximal and distal convoluted tubules staining. Scale bar = 50 μm. g. Region with T Cells (CD3 and CD4 positive) Insets highlight overlapping staining between CD3 and CD4. Scale bar = 20 μm.

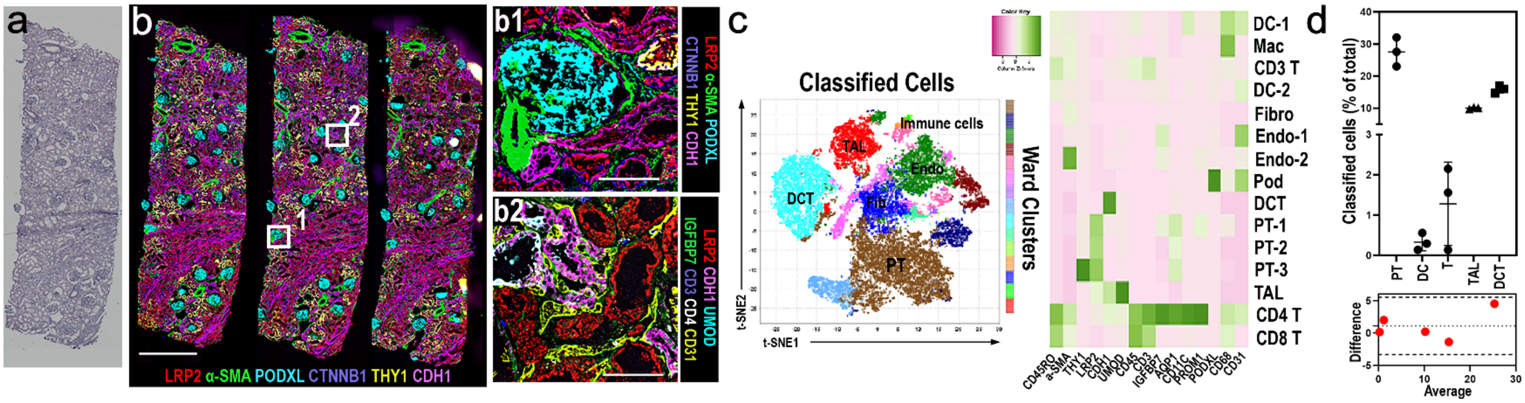


**Supplemental Figure 2: Codex reproducibility**

(a) H&E staining of a biopsy-size kidney section after CODEX imaging (b). (b) CODEX imaging of consecutive (1,2) or alternating (2,3) sections labeled with 22 markers had comparable staining patterns (scale = 1 mm). Boxes a and b, enlarged at right, stained with markers for glomeruli (PODXL), tubules (LRP2, UMOD) and immune cells (CD3, CD4) have expected staining patterns (scale = 100µm). (c) Unsupervised clustering with VTEA identifies major cell types/subtypes based on marker expression. d) The proportions of classified cell types are similar for the three sections in b and a pairwise comparison demonstrates a reproducible pipeline (bottom).

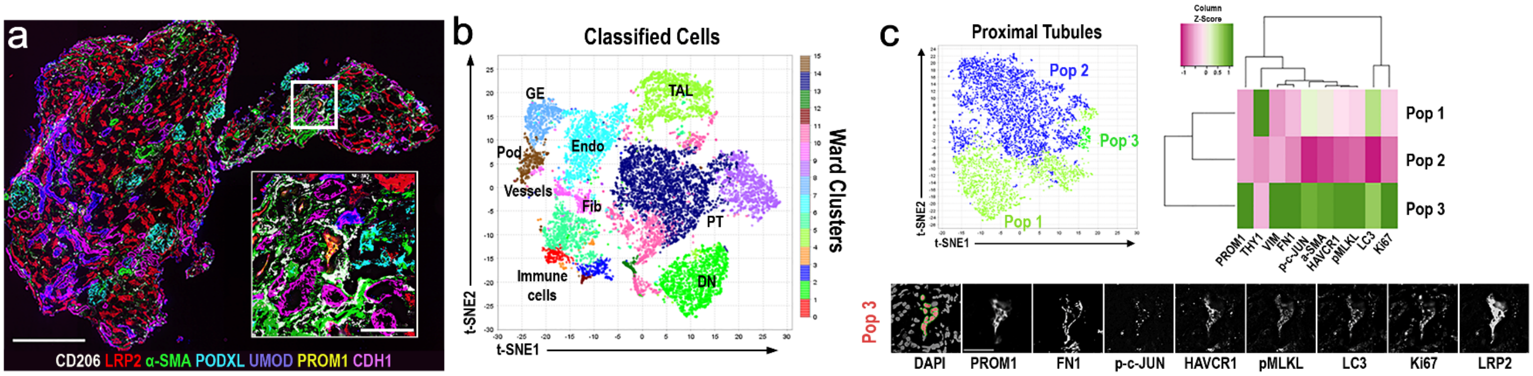


**Supplemental Figure 3: Validation of injury profile of proximal tubules using extended CODEX panel.**

(a) CODEX imaging of a cortical biopsy section using 38 markers; scale = 500µm. Boxed area is enlarged in inset (scale = 100µm). (b) Unsupervised analysis using VTEA and identification of major cell types. (c) Proximal tubule (PT) cells were re-clustered identifying 3 distinct populations (Pop) based on the expression of injury/ proliferation/ fibrosis markers. Pop1 was uniquely THY1-positive, and Pop3 was uniquely PROM1-positive and expressed several injury markers. Mapping back of Pop3 cells (bottom, red nuclear overlays) in the image volume showing the staining of the injury and fibrosis markers used, validating the unsupervised classification. (scale = 50um)

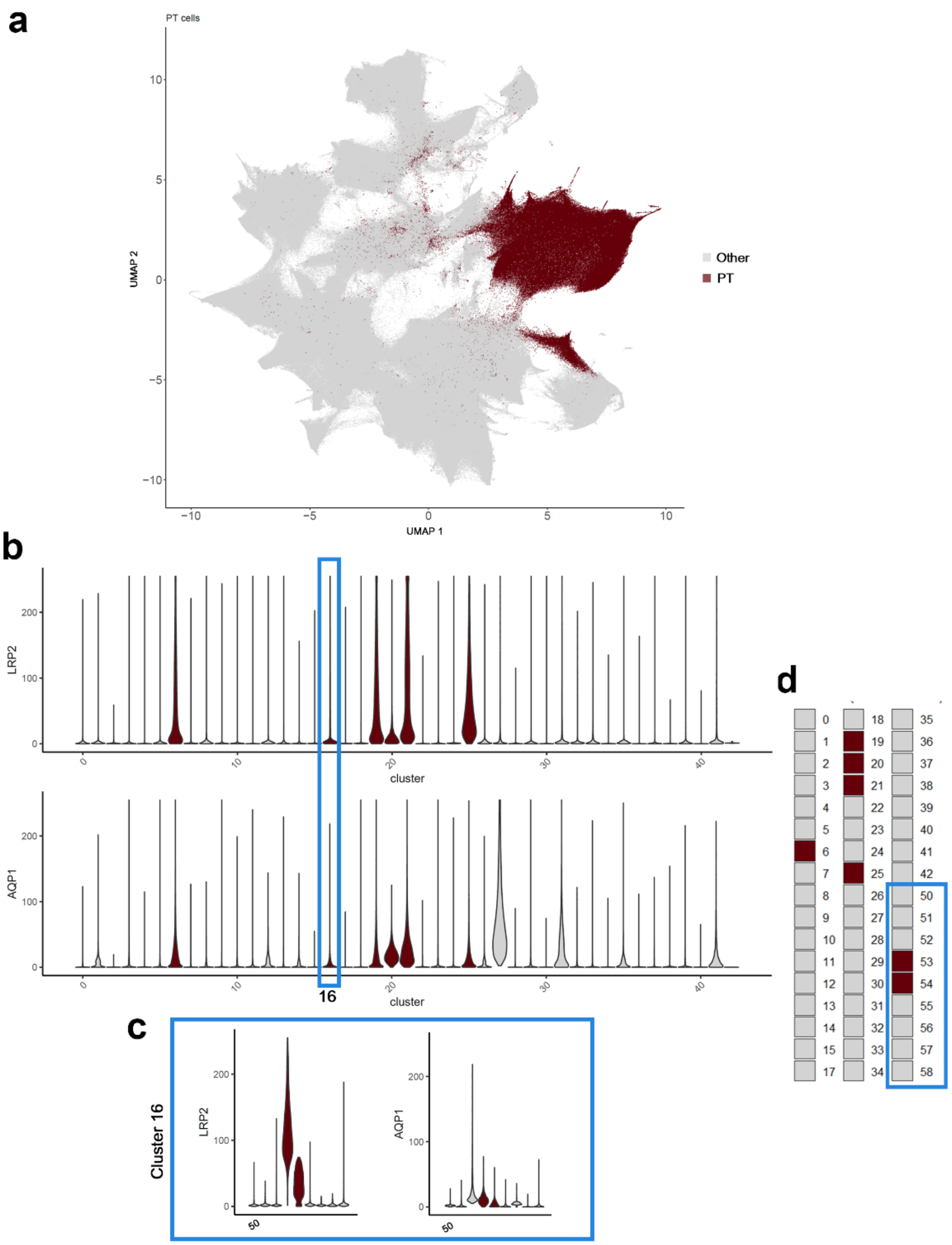


**Supplemental Figure 4: Identification of proximal tubules in the Phenocycler validation dataset based on LRP2 and AQP1 expression.**

**(**a) UMAP feature plot showing all proximal tubular (PT) cells (441,509 cells in brown color) in the entire dataset comprising ~ 2.35 million cells. (b) Violin plots depicting the expression of LRP2 and AQP1 for the 41 clusters which were generated using Louvain algorithm (see methods) highlighting PT clusters in brown color. Cluster validation was done with back-mapping on imaging data, and identified clusters 6, 19, 21 and 25 are pure PTs, whereas cluster 16 was mixed. (c) sub-clustering of cluster 16 further identified sub-clusters 53 and 54 as pure PTs. (d) Representation of PT clusters (6, 19, 10, 21 & 25) in the parent Louvain clusters (0-15 & 17-42) and sub-clusters (53 &54) in parent cluster 16 comprising all the PTs shown in (a).

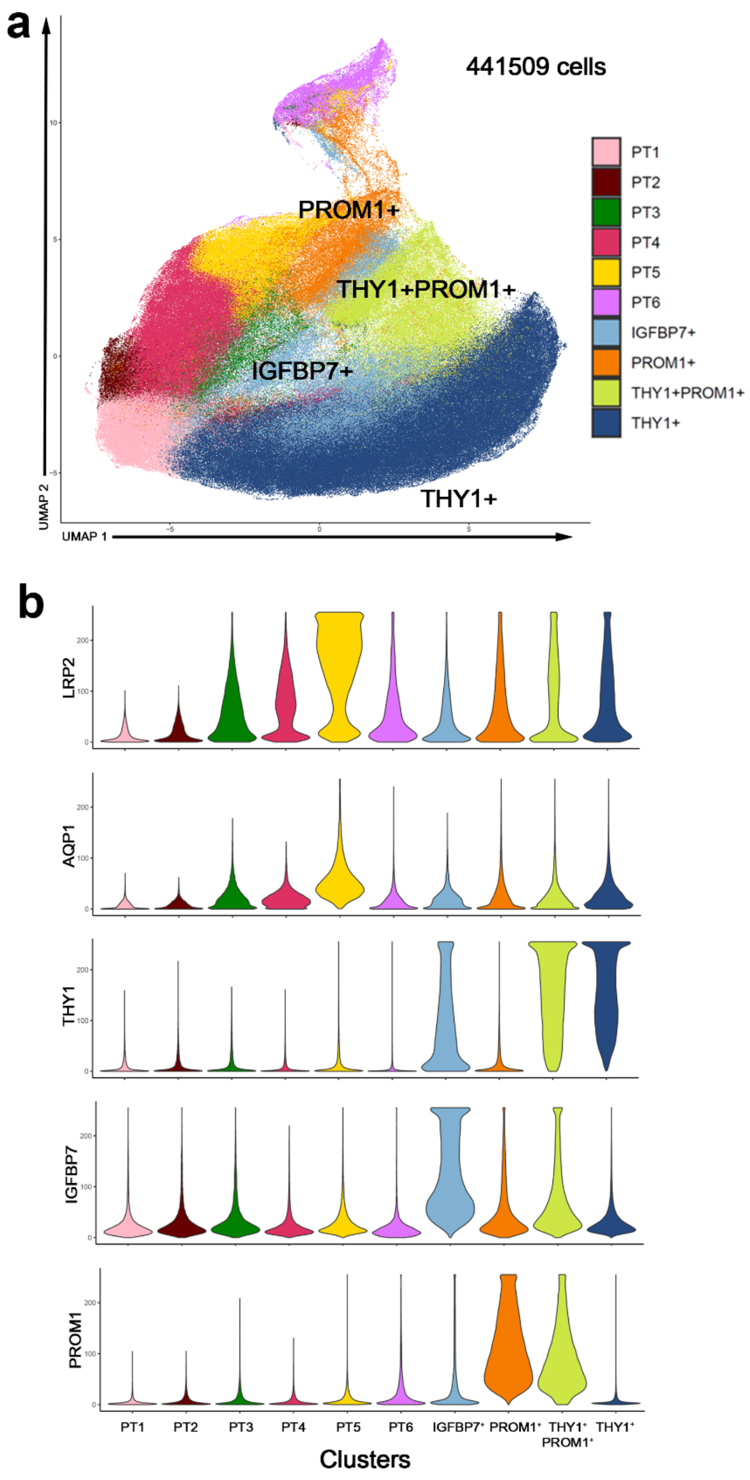


**Supplemental Figure 5: Proximal tubule sub-clustering in the Phenocycler validation dataset.**

(a) UMAP showing the PT clusters that were grouped based on THY1, PROM1 and IGFBP7 expression (see methods). (b) violin plots for the expression of various markers for the grouped PT clusters.

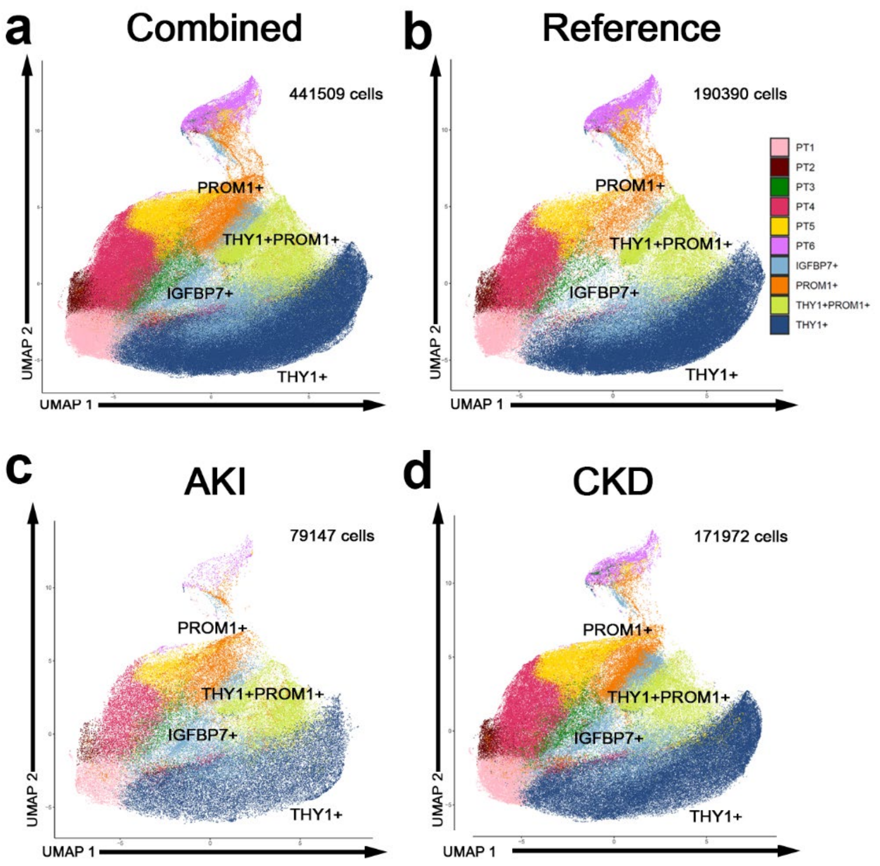


**Supplemental figure 6: Integrated PT clusters by condition in the phenocycler validation dataset.**

UMAP embeddings of the PT clusters from various conditions: (a) combined PTs (b-d) PTs from reference, acute kidney injury (AKI), and chronic kidney disease (CKD), respectively.

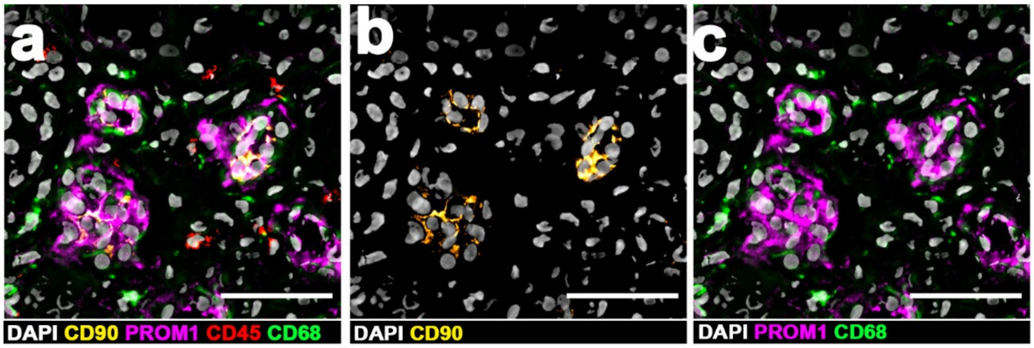


**Supplemental Figure 7. CODEX imaging of Proximal Tubule Epithelium in reference tissue**.

(a) subregion of reference tissue and select epithelial cell markers. (a) Epithelial cells are labeled with DAPI(Gray), CD90/Thy1 (Yellow), PROM1(Cyan) or CD68(green). (b & c) mixed populations of CD90/Thy1, CD68 and PROM1 positive epithelium. Scale bars = 50 µm

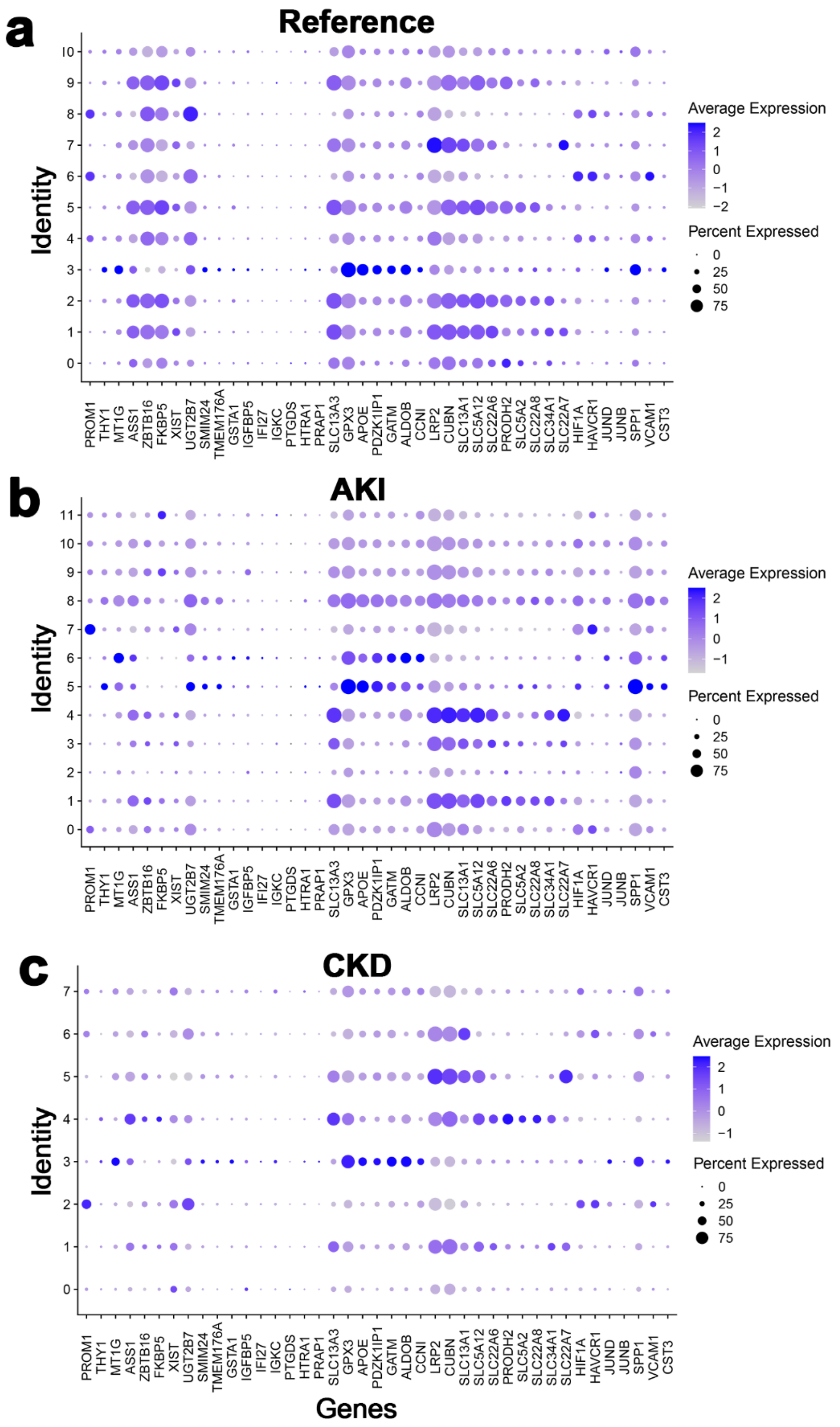


**Supplemental Figure 8: Single nuclear transcriptomic signatures of various PT subpopulation based on condition**

PT clusters from reference (a), acute kidney injury- AKI (b) and chronic kidney disease CKD (d) from the KPMP snRNAseq atlas displaying genes of interest (upregulated genes in THY1+ PT cells from Figure 3 and canonical PT and injury genes).
